# Drinking earth for wine. Estimation of soil erosion in the Prosecco DOCG area (NE Italy), toward a soil footprint of bottled sparkling wine production

**DOI:** 10.1101/516245

**Authors:** SE Pappalardo, L Gislimberti, F Ferrarese, A Garlato, IC Vinci, M De Marchi, P Mozzi

## Abstract

Prosecco, one of the most widespread sparkling wine in the world, is produced in Northeast Italy by a rate of 400 M bottles per year, with the fastest growing demand in the global market at present. A production of 90 M bottles year^−1^ is currently running in the historical Prosecco sector (215 km^2^), defined as the Controlled and Guaranteed Designation of Origin (DOCG) area, in a steep hilly landscape of Veneto Region (Conegliano-Valdobbiadene) registered in 2017 for the UNESCO World Heritage tentative list. To sustain wine production agricultural intensification boosted to re-setting of hillslopes and land use changes toward new vineyard plantations. The aim of this study is to assess soil erosion rate, calculating a sort of “soil footprint” for wine production by i) estimation of the total soil erosion, ii) identification of the most critical areas, iii) simulation of different nature-based mitigation scenarios. RUSLE model was adopted to estimate soil erosion in Mg ha^−1^ year^−1^, using high resolution topographic data (LiDAR), 10 years rainfall data analysis, detailed land use and local soil characteristics.

We found that the total soil erosion estimation for the Prosecco DOCG area is 546,263 Mg year^−1^, with an erosion rate of 25.4 t ha year^−1^, which is 11 times higher than the Italian average. Prosecco vineyards contributes to 400,000 Mg year^−1^, by a mean rate of 59.8 Mg ha^−1^ year^−1^, and encompass 74% of all the erosion in the whole DOCG area. Soil erosion modelled is mainly concentrated in cultivated hillslopes, highlighting critical areas with more than 40 Mg ha^−1^ year^−1^), mainly clustered on steep slopes.

The modelled soil loss of a single bottle of Prosecco is, therefore, about 4.4 kg year^−1^. In contrast, alternative scenarios of different nature-based mitigation measures (hedgerows, buffer strips, and grass cover) showed that total erosion in the Prosecco DOCG area would be reduced to 275,140 Mg year^−1^, saving about the 50% of soil. In vineyards a general decrease of almost 3 times (from 400,000 to 135,161 Mg year^−1^) is also demonstrated, reducing on average the erosion rate from 59.8 to 19.2 Mg ha^−1^ year^−1^. This study highlights, thus, that an integrated soil erosion monitor system is needed in the DOCG area as well as the implementation of nature-based mitigation measures as sustainable agricultural layout for modern agroecosystems.

## Introduction

### Agricultural lands and soil erosion

Agricultural lands presently occupy about 37.4% (56.1 M km^2^) of the 150 M km^2^ of Earth land surfaces [1]. They amount to the 50% if glaciers, deserts, rocks, and other physical environments not suitable for agriculture are excluded [2–4]. Indeed, agricultural lands are the widest Human-modified ecosystems, making crop production the most extensive form of land use on Earth [5]. The geographical dimension at global scale of agriculture is crucial to understand the role it plays in terms of land degradation and erosion processes, which are boosted up to 1–2 orders of magnitude greater than the natural rates of soil production [6]. In fact, high erosion rate in conventional farming are mainly linked to unsustainable soil management and agricultural practices: intense tillage, soil compaction due to the use of heavy machinery, down-slope cropping on hillside, and intensive herbicide application [7,8]. Recently, it has been estimated that soil erosion directly linked to mismanagement of agricultural lands affects about 5,520,000 km^2^ worldwide [7]. As results of heavy soil erosion, about 30% of the world’s arable land have been already lost and turned to unproductive [9].

In Europe 12.7% of total land surface is affected by moderate to high soil erosion risk [10]. This means that a total area of about 14 M ha (a surface wider than Greece), loses soil at a rate of 2.46 Mg ha^−1^ year^−1^ on average, resulting in a total annual soil loss of 970 M Mg [11,12]. According to estimation based on erosion plot data, the mean erosion rate of total surface in Italy is 2.3 Mg ha^−1^ year^−1^, which represents the 12.5% of the total European erosion [13]. Due to unsustainable agricultural practices of intensive crop production, soil erosion is one of the main environmental concern in many sectors of Southern Europe, especially in sloping rainfed croplands. Many field-based researches performed in Spain demonstrated that agricultural practices based on herbicides and conventional tillage results in high erosion rates: Gomez et al. (2003) found that on slopes up to 20% soil erosion could reach 80 Mg ha^−1^ yr^−1^ [14]; Ramos et al. (2008) measured soil pro?le lowering due to particle detachment of up to 0.2 ± 0.1 m yr^−1^ along slopes ranging from 2 to 45% in an orchard conventionally tilled [8,15]; Keestra et al. (2016), by means of simulated rainfall experiments and soil analyses in apricot orchards, demonstrate that tillage and herbicide treatments should be avoided to control soil erosion [16]; Cerdà et al. (2009) found that soil erosion rates in citrus orchards plantations were 2 Mg ha^−1^ after 1 hour of a 5-year return period rainfall thunderstorm [17].

Among agricultural lands, vineyards cover about 76,000 km^2^ of the Earth surface, an area wider than Ireland, mainly oriented to wine production[18,19]. However, about half of world vineyards surfaces is cultivated in Europe (33,000 km^2^) whose 30% is mainly concentrated in Italy (6,950 km^2^), Spain (9,670 km^2^), and France (7,870 km^2^). Vineyards are respectively 2.3%, 1.9%, and 1.2% of the country area [18,19]. Aside from representing one of the most important cultivations in terms of local economies, income, and employment, vineyards recently gained an increasing attention since it is one of agricultural land use that causes the highest soil erosion rates [20–23].

### Soil erosion in Mediterranean vineyards

Due to geomorphological, climatic, and edaphic conditions together with anthropogenic factors vineyards in Mediterranean ecosystems are particularly inclined to land degradation and soil erosion [20,24]. Agricultural lands for vineyards are often located on hilly areas, on steep slopes, resulting in the highest measured soil erosion compared to rainfed cereals, olives groves plantations or scrublands [24]. In fact, topography is one of the dominant factor affecting soil erosion and sediment transportation. In addition, Mediterranean vineyards have to face high intensity rainfall events, mainly concentrated in Spring and Autumn. As well documented, soil erosion processes are strongly influenced by the high magnitude – low frequency rainfall events which presently have to be even more considered in the climate change scenarios [24,25]. Furthermore, Mediterranean lands are generally poor in nutrient and organic matter content which are key factors on soil stability and erodibility [26]. Finally, the market-driven farming intensification of Mediterranean vineyards for wine production results in unsustainable soil management: common practices are mainly based on deep mechanical tillage and chemical weeding without tillage. Both soil management systems result in bare soil during most of the year, leaving wide areas exposed to the rainfall, with a notable increase in runoff and soil erosion rate [27,28]. Different soil management systems and agricultural practices result in measured soil erosion rates which range from 3.3 to 161.9 Mg ha^−1^ yr^−1^ [24,29].

### Prosecco DOCG

The international wine trade in 15 years grew by 75% in volume and doubled in value, leading in 2015 to a total volume of import equal to 98 million hectoliters. Considering the last five years, with the exception of Champagne, sparkling wine continued to grow with an annual rate of 7% in value and 6% in volume, turning the Prosecco to an emblematic case as one of the most exported in the world [23,30]. Specifically, Prosecco wine production boosted from 2009 after the “Protected Designation of Origin” (PDO) by labelling the Controlled Denomination of Origin (DOC), and the Controlled and Guaranteed Denomination of Origin (DOCG) areas to identify two specific growing areas. In the last decades the Prosecco wine production has notably increased in the DOCG area due to a combination of global market demand and large investments in the region which boosted both crop production and land use change into vineyards croplands [31–33]. In 2017, the Prosecco DOCG growing area was officially enrolled in the tentative list the for the UNESCO World Heritage status [23,34]. However, the UNESCO candidacy was criticized both at academic and at civil society levels due weaknesses in terms of the socio-environmental unsustainability of Prosecco farming system [32]. The dispute was fueled since in July 2018 the 42^nd^ World Heritage Committee rejected as first evaluation the Prosecco candidacy because it does not meet the UNESCO criteria [35,36].

The Prosecco DOCG vineyards increased from some 4,000 ha in 2000 to 5,700 ha in 2010, and beyond 7,000 ha officially declared in 2016 [34,37,38]. At present, Prosecco wine production is over 400 M and 90 M bottles respectively in the DOC and DOCG geographical areas [39]. In such context the economic and production factors are driving drastic changes in land use, undermining an ecosystem stability based on soil system, and fueling the debate about the sustainability of vineyards cropland.

Considering the complexity of the phenomenon and its implications at socio-economic and environmental level, therefore there is an urgent need to assess and to estimate the amount and the potential rate of soil erosion at agricultural landscape scale. Furthermore, modelled erosion rate at a very detailed scale and simulated nature-based scenarios would represent a scientific contribute to support and design a more sustainable land management for wine production, especially in sensitive areas where soil erosion is over the tolerable threshold.

The general aim of our study is to assess soil erosion at landscape scale in the Prosecco DOCG growing area, calculating a sort of “soil footprint” for bottled wine production. Specific aims are: i) to quantify the total soil erosion; ii) to identify the most critical areas in term of soil erosion rates; iii) to simulate alternative sustainable scenarios to reduce soil erosion processes and off-site impacts, applying possible mitigation measures at field scale.

## Physical geography of Prosecco DOCG

The study area falls within the Prosecco DOCG wine production area which spans 215 km^2^ in the North-East sector of Italy (Province of Treviso) and it encompasses fifteen small-medium Municipalities, in a scattered urban-agricultural territorial matrix. Vineyard cropland presently occupies the 32% of the DOCG area, representing one of most diffuse cultivation (Fig 1a).

**Fig 1a.**
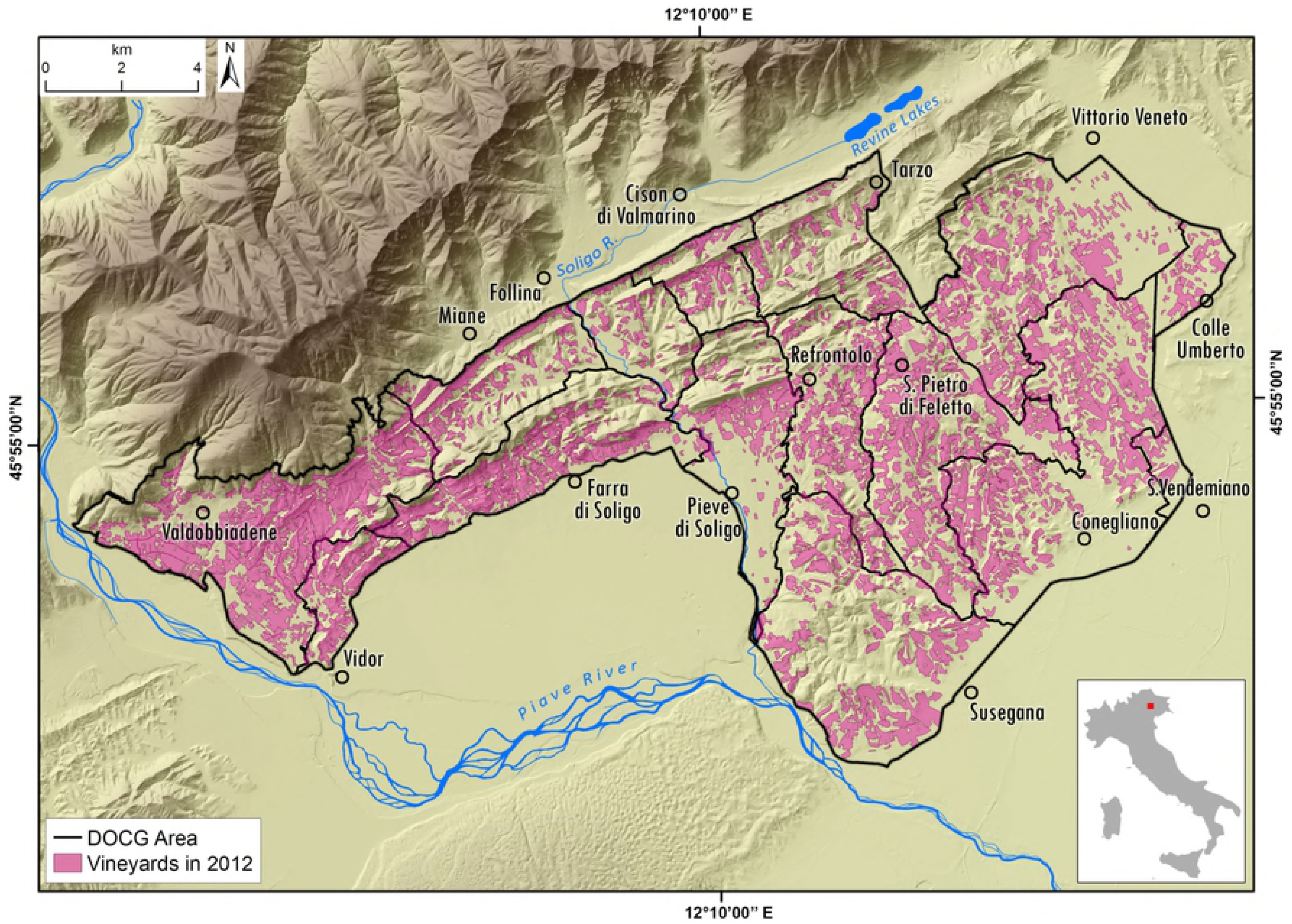
Geographical and geomorphological setting of the Prosecco DOCG area. **Fig 1b.** Vineyards distribution in the Prosecco DOCG area.

Generally, Prosecco vineyards concentrate in the south-facing slopes, while copses and chestnuts are in the north-facing ones. In the hilly region, modifications in geomorphology and, therefore, changes in the drainage systems, are often related to crop production intensification and to the high levels of mechanization and standardization required; hence, modern hydraulic-agrarian layouts by vertical ploughing with vineyard rows setup along the steepest slope are now generally preferred. On the contrary, contour farming by traditional or modern agricultural terraces are limited, and have been substantially reduced in the past years. At field scale, about 30-60% of grass cover is generally maintained between vineyards rows.

The landscape has elevation ranging from 60 to 500 m a.s.l., and it is principally dominated by 70% of hilly terrain, and 28% of alluvial plain, while only 2% is mountainous (Fig 1b). The wide and fragmented agricultural landscape is currently dominated by intensive Prosecco cropland (about 86% of the whole cropping system - Figs 1b and 2) which is extended both in the upper alluvial plain and in the hilly areas, which are often scarcely accessible and have steep slopes.

**Fig 2.**
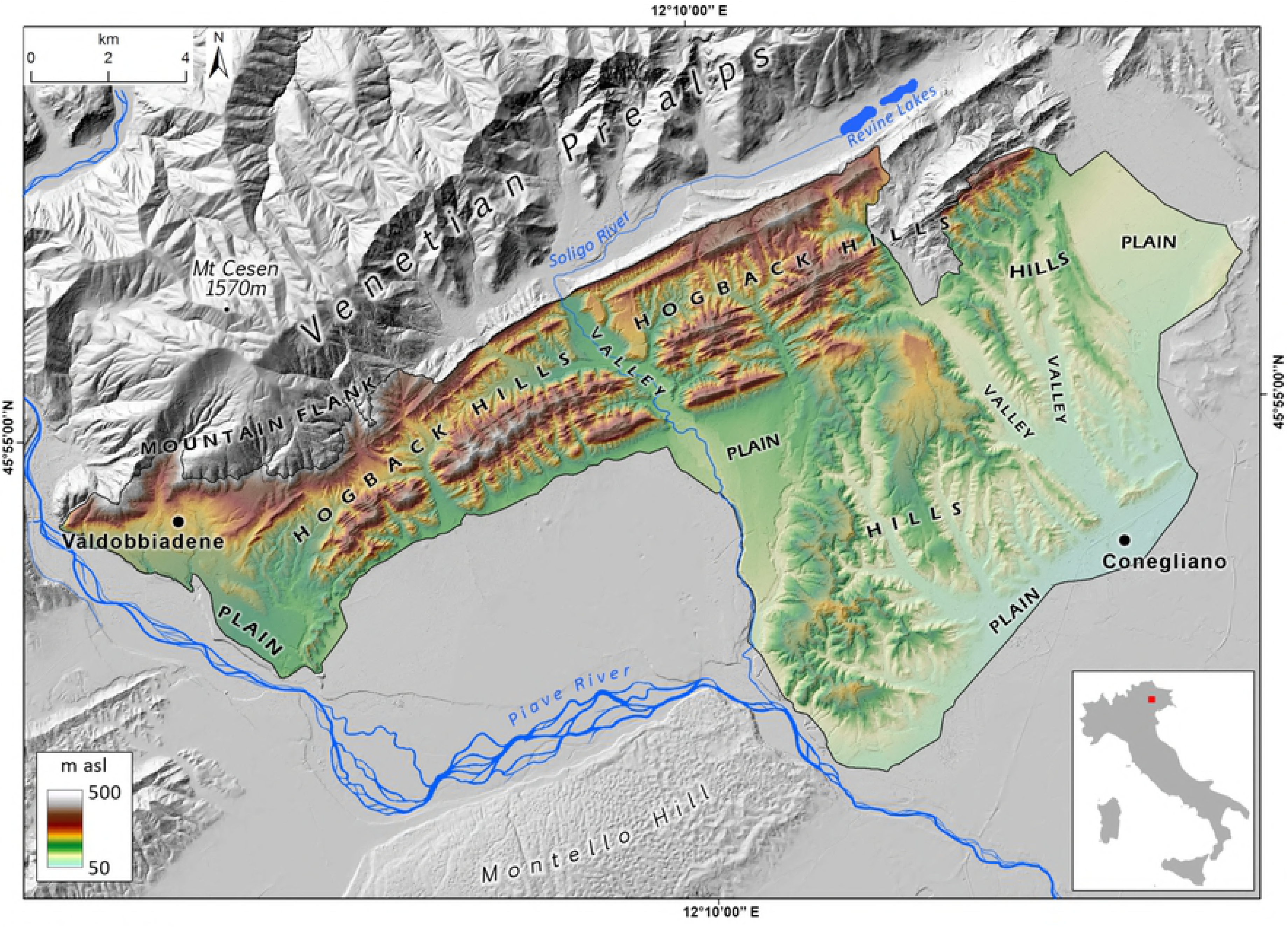
Percentage of area covered by principal land use classes in the Prosecco DOCG zone. More of 30% of territory is covered by vineyards.

According to Köppen climate classification, the Prosecco DOCG area is at the transition between a temperate oceanic climate (Cfb) and a Mediterranean type with hot dry-summer (Csa). According to Thornthwaite (1948) climate classification is B3 Humid (60-80 moisture index). Mean precipitation is 1,200 mm year^−1^ and average temperature is 12.7° C.

Geology of the area relates to the collision between the Adria micro-plate to the South and the Euro-Asiatic plate to the North during the Cenozoic, which produced the uplift of the Italian Southern Alps [40]. In the southernmost Alpine sector the crustal thickening is equilibrated by a series of parallel, south verging thrusts with direction NE-SW (Bassano-Valdobbiadene thrust, Valsugana thrust, Montello thrust) [41]. The Prosecco DOCG area lies in a zone between two parallel thrusts: the Montello and the Bassano-Valdobbiadene thrust. The stratigraphic succession that is cropping out in the study area spans from Mesozoic dolostone and limestone to Upper Miocene conglomerates (Conglomerato del Montello), sandstone and marls [42].

The Montello thrust and its ramp anticline contributed to the formation of the Quartier del Piave intermontane valley, instead the “Refrontolo syncline” generated highly inclined strata that are shaped as hogback hills. The geomorphologic landscape of the study area is strongly shaped by this series of long, NE-SW oriented ridges, ranging in elevation between 50 and 500 m a.s.l. These landforms are typical hogbacks, modeled in alternances of conglomerate, marls and sandstone [42]. The alluvial plain sector in the Quartier del Piave is characterized by predominant gravel deposits, deposited since the Last Glaciation to present by the Piave River, the Soligo River and other minor streams [42,43].

Fourteen landscape-soil units characterize the study [44]: fans, alluvial terraces and valley fills by Prealpine streams of the Last Glaciation with leached soils (C1), and of Holocene age with poorly developed soils (C2); gravelly plain of the Piave River with leached, carbonate-depleted and rubified soils (P1), leached and carbonate-depleted soils (P2), and poorly developed soils (P6); fine-grained alluvial plain of the Monticano and Meschio Rivers, with poorly developed soils (M3); terminal moraines of the Last Glacial Maximum (LGM) with slight evidence of carbonate leaching (G2), and of pre-LGM glaciations with leached and rubified soils (G1); steep hillslopes in conglomerate, with shallow and poorly developed soils (H1); low-gradient hillslopes in conglomerate, with strongly decarbonated, rubified soils with evidence of clay illuviation (H2); steep hillslopes in sandstone, with moderately deep and developed soils (H3); low-gradient hillslopes in marls and siltite, with moderately deep and developed soils (H4); long and steep mountain slopes in massive and hard limestone, with shallow and poorly developed soils (V1); long and steep mountain slopes in well-stratified, moderately resistant limestone, with moderately deep and leached soils with clay illuviation (V2) [44].

## Material and methods

### RUSLE model

Different models and field-based approaches were developed to assess spatial distribution of soil erosion. Among them, the use of empirical models combined with spatial data processed into Geographical Information Systems (GIS) is the widest tool to quantitatively estimate and map soil erosion rates. To estimate soil erosion in the study area, we adopted the Revised Universal Loss Equation (RUSLE) defined by Renard *et al.* [45] and derived from the Universal Soil Loss Equation (USLE), previously proposed by Wischmeier and Smith (1978) [46]. RUSLE is the most widely-used empirical model for soil erosion estimation at landscape scale [9,12,47,48]. It was also tested in several study cases in Mediterranean context, both at basin and landscape scale [22,49–51]. RUSLE model is based on the main factors which strongly contribute to soil erosion processes, combining data about topography, soils, rainfall, and land use in a GIS environment. It performs a spatial simulation of the erosion processes quantifying soil loss in terms of Mg ha^−1^ year^−1^. According to quality and geometric resolution of spatial data, by running the RUSLE model it is possible to identify the magnitude of soil erosion processes at landscape scale and map it [9]. The RUSLE model is based on the equation:

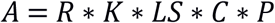

The RUSLE model is based on five independent variables: i) R, ii) K factor, C, iii) LS factor; iv) C factor; v) P factor.

To perform RUSLE model we collected and modelled spatial and temporal data for each factor: i) meteorological data based on 20 local weather stations (R factor) (Fig. 3); ii) pedological data about the erodibility of soils and its susceptibility to erosion (K factor); high resolution topographic data (LS factor); and land use data at regional scale (C factor).

**Fig. 3.**
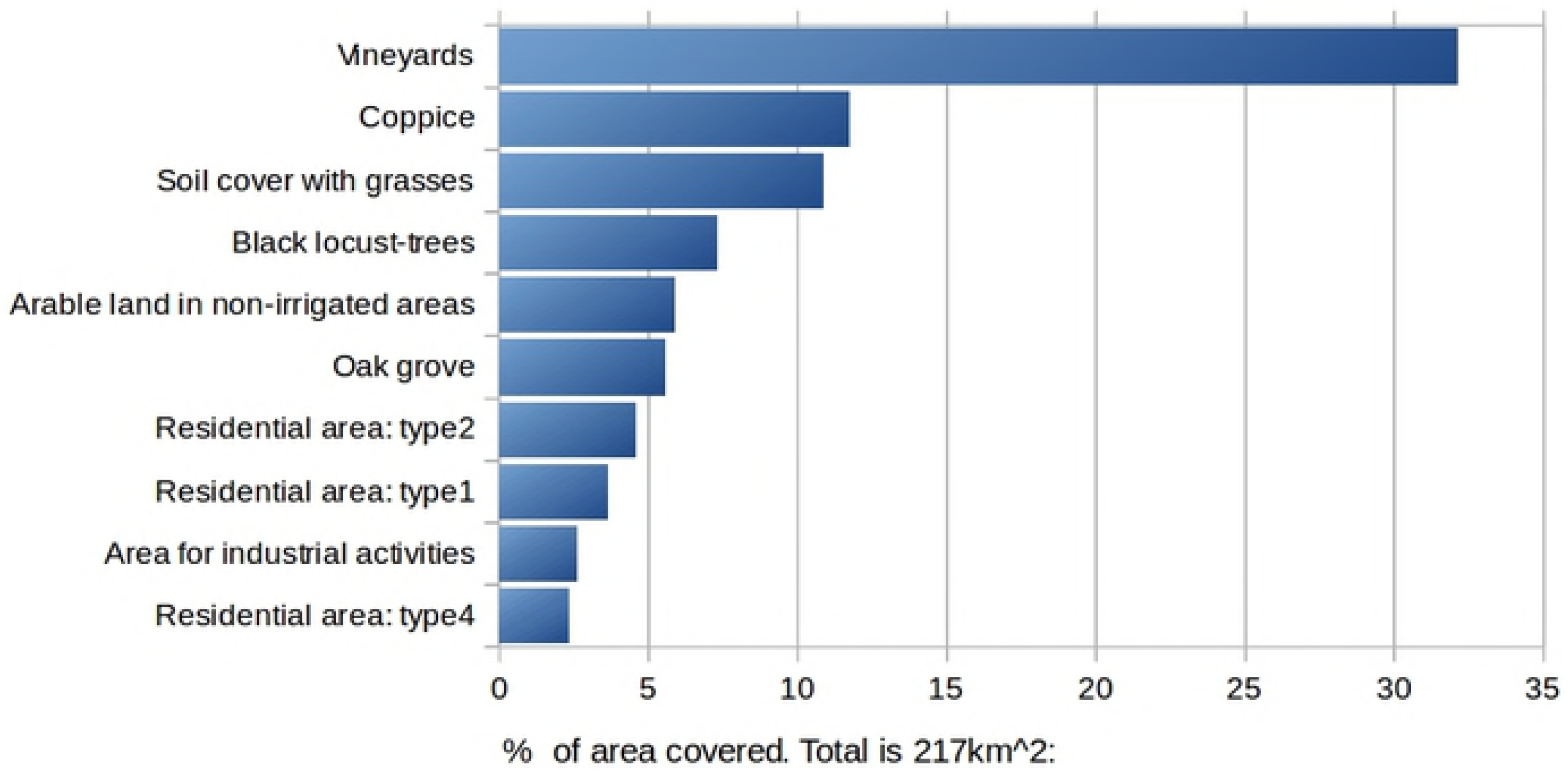

### R factor

R factor represents the energy and ability of the rainfall to erode soil. It is strictly related to the main impulsive rainfall events for a specific region [52]. According to local climatic trends, the availability and the spatial distribution of meteorological data, different empirical formulas to calculate R factor were developed by several researchers. Climatic data from the Regional Agency for Environmental Protection and Prevention of the Veneto (ARPAV, 2012), shows for the Veneto Region typical rainfall of modest intensity distributed during the year in two main peaks: one during autumn and another one in spring. Hence, according to local climatic conditions we used the formula that represents the best fit for such regime, according to Wishmeier and Smith (1987) and revision by Renard *et al.* [45]:

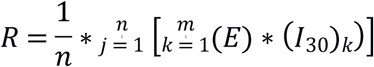

Where:

*n* represents the number of years considered, that in this study are from 2006 to 2015;

*k* is the rainfall number of half hour events;

*E* is the rainfall energy event estimation;

*I*_*30*_ is the amount of rain in 30 minutes;

*m* is the summarize of every rainfall events for all considered years.

*E* is calculated using the following expression:

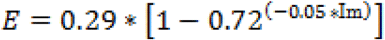

where *Im* is the intensity for 5 minutes of rainfall recorded by 20 local weather stations of ARPAV, distributed on all Region.

To calculate R values for each of the 20 weather stations we wrote a specific algorithm and performed it by using R software (R Core Team, 2016) performing a rainfall analysis on 10 years of time-series. The mean R values for each weather stations were, therefore, spatially interpolated using the Inverse Distance Weighted (IDW) algorithm in GIS environment.

### C factor

C factor defines the type of soil cover that influences soil erodibility. Vegetation or artificial cover reduce the erosion effect. The anthropic pavements are impermeable and immobilize the soil. Vegetation has a double effect: leaves partly intercept rain drops, lowering the rainfall kinetic energy at impact with ground (“spalsh erosion”), roots promote water infiltration in the topsoil, lowering surface runoff.

To calculate the C factor, we used the IV Level of CORINE-based dataset at regional scale [53,54]. There are many studies that use several methods to determinate a suitable C value for different type of land use in different morphoclimatic conditions. We therefore selected C values found in literature that best fit the regional environmental conditions [55,56]. The C factor value for each type of land use is traditionally defined by numerous empirical equations and sample points that consider vegetation characteristics and morphological conditions, as surface roughness and surface cover [45]. Concerning vineyards there are different values for C factor in literature (from 1 for arable land, to 0.02 of olive groves), according to the hydraulic-agrarian layouts, percentage of grass cover between rows, and the different techniques and crop practices adopted [57]. This happens because the viticulture is distributed in different climatic zones, from rainy alpine valleys to semi-desert regions in Mediterranean country. In our study, to calculate RUSLE index we adopted a conservative value of 0.12 for vineyards land use as suggested by ARPAV (2008) [55].

### LS factor

LS factor represents the topography (length and slope) that influences soil erosion effectiveness [58]. This factor indicates where erosion may act more aggressively and it includes only topographic variables. Depending on these morphological conditions, especially after intense rainfall events, water can acquire high velocity and energy in order to form streams or erosional channels. In order to calculate LS factor, we performed a DTM analysis over 1 m geometric resolution of Laser Imaging Detection and Ranging data (LiDAR) [59]. The formula used for LS factor estimation is after Moore and Burch (1986):

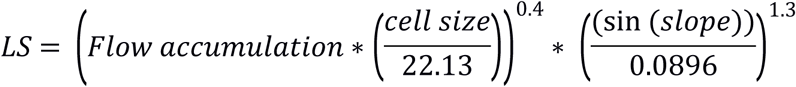

where *Flow accumulation* is calculated for each pixel, as the sum of area that lies upstream of the respective basin. In this work the cell size adopted is 1 m. Slope raster was built by function “terrain” in R raster library (R Core Team, 2016) and represents the slope in degrees.

*Flow accumulation* is obviously very high in correspondence of rivers and streams, which results in high LS values that have an important impact on RUSLE final calculation. As our study focuses on areal soil erosion related to surface runoff and incipient rills and gullies, rather than linear erosion along the river network, we chose to exclude watersheds wider than 5 ha from the calculation of flow accumulation.

### K factor

K factor represents the erodibility of soils and its susceptibility to erosion. The method used for calculation is the original equation of Wischmeier, described in 1978, as reported in Handbook 703 “Predicting soil erosion by water: a guide to conservation planning with the Revised Universal Soil Loss Equation” [45].

In literature, due to the difficulty in recovering soil data required by the original equation, simplified pedofunctions are available for calculating the K factor. At regional level the K factor was calculated using Renard and Torri simplified function (1997) [45,60]: the former requires only values of sand and silt, the latter requires also organic matter. By comparison of results, it was preferred to use data derived from Wischmeier (1978) [46], that considers various soil characteristics, as shown below:

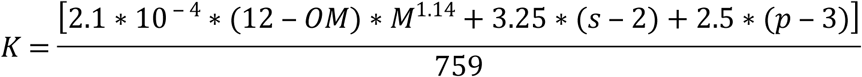

where:

OM: percentage of organic matter in topsoil;

M: textural parameter (depending on sand, fine sand and silt percentage);

s: structure class code;

p: profile permeability class code.

All soil data were available in ARPAV database; the K factor was calculated for each type of soil (soil typological unit) and then spatialized relying on the 1:50,000 soil map, available for the whole DOCG area [44].

## RUSLE analysis

All the data were collected and assembled together to perform RUSLE spatial analysis in Prosecco DOCG area, at 1 m pixel^−1^ geometric resolution. We analyzed the soil erosion in terms of magnitude (Mg ha^−1^ year^−1^) and mapped it by using GIS Zonal Statistics tools. By running the RUSLE model we estimated which areas show high values of soil erosion and where they are located. We reclassified RUSLE output values in 4 classes: low value (0-4 Mg ha^−1^ year^−1^), medium value (4-10 Mg ha^−1^ year^−1^), high value (10-40 Mg ha^−1^ year^1^), very high value (more of 40 Mg ha^−1^ year^−1^). By RUSLE analysis we evaluated land use influence on soil erosion phenomenon. We calculated the soil loss per hectare (Mg ha^−1^ year^−1^) and the total loss (Mg year^−1^). We evaluated soil erosion potential in the landscape, classifying RUSLE values on regional soil units. We also evaluate the soil loss at Municipality scale in order to highlight the wine-producing district most exposed to erosion processes. Moreover, we used the RUSLE model results to calculate a sort of “soil footprint” for wine bottles using official production data of Prosecco DOCG published in 2017 [39].

### Soil erosion under alternative nature-based scenarios

Within the CAP framework, EU promoted the adoption of “best practices” in soil management to control erosion processes by keeping the land under “Good Agricultural and Environmental Condition” (GAEC). Different landscapes features such as grass cover, dry-stone walls, reverse-slope benches on one side, and hedgerows or buffer strips to reduce runoff volume and protect habitats, are included in GAEC standards [12,61]. We therefore performed four different scenario simulations at Prosecco DOCG scale, by adopting four different nature-based mitigation measures to increase agricultural sustainability and to protect surface water from loose of herbicides and pesticides: hedgerows, grassed buffer, and a grass cover between inter-rows of vineyards.

In scenario 1 we assigned a conventional grass (C factor 0.005) buffer zone, of 5 m from tail lift of rivers and streams with a minimum value of 2^nd^ order; in scenario 2: we assigned 3.5-m hedgerows of shrub (C factor 0,003) as buffer zones around the vineyards; in scenario 3 we modeled a combined scenario summarizing the effects of scenario 1 and scenario 2. Finally, in the fourth one, we simulate the most sustainable agricultural best practices scenario without the application of herbicides in land management: we simulate to keep grass cover in 100% of vine inter-rows during the Winter period. According to Bazzoffi et al. (2017) we used 0.04 value C cover for grassed inter-row vineyard management [56]

## Results

### Soil erosion estimation: actual scenario

RUSLE analysis showed that the total soil erosion estimation for the Prosecco DOCG area is 546,263 Mg year^−1^, by a rate of 25.4 Mg ha year^−1^ on average. Beyond this, more of 70% of the total surface showed a potential soil erosion between 0 and 4 Mg ha^−1^ year^−1^, 12% is between 10 and 40 Mg ha^−1^ year^−1^, while the 12% is more 40 Mg ha^−1^ year^−1^ (Fig 4a).

**Figure 4A:**
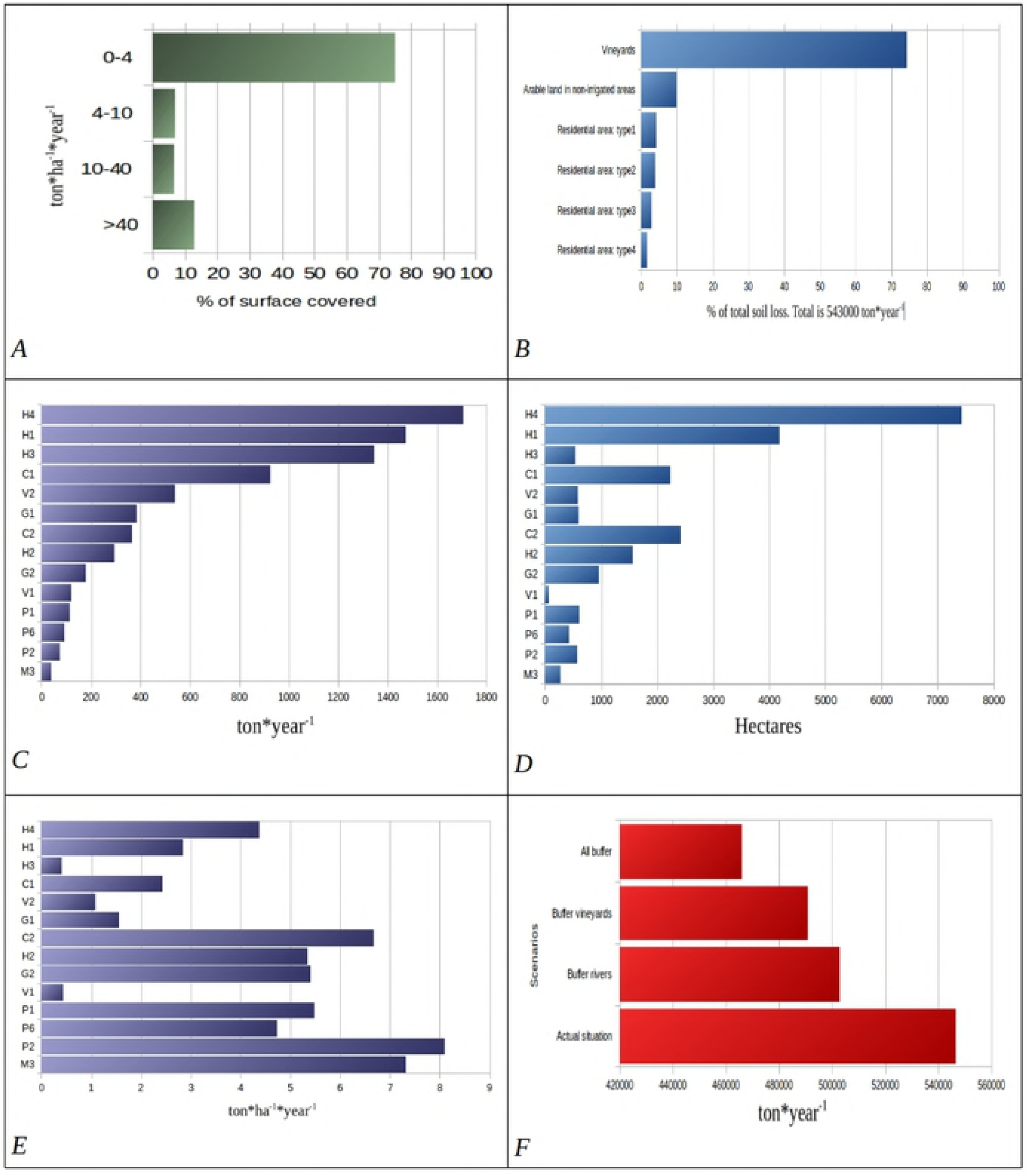
Percentage of the area in RUSLE erosion classes: low erosion (0-4 Mg ha^−1^ year^−1^), medium erosion (4-10 Mg ha^−1^ year^−1^), high erosion (10-40 Mg ha^−1^ year^−1^) and very high erosion (>40 Mg ha^−1^ year^−1^). **Figure 4B**: Percentage of potential soil loss from RUSLE modelling in different landuse. **Figure 4C, 4D and 4E:** Soil erosion along the landscapes-soil units. Fans, alluvial terraces and valley fills by Prealpine streams of the Last Glaciation with leached soils (C1), and of Holocene age with poorly developed soils (C2); gravelly plain of the Piave River with leached, carbonate-depleted and rubified soils (P1), leached and carbonate-depleted soils (P2), and poorly developed soils (P6); fine-grained alluvial plain of the Monticano and Meschio Rivers, with poorly developed soils (M3); terminal moraines of the Last Glacial Maximum (LGM) with slight evidence of carbonate leaching (G2), and of pre-LGM glaciations with leached and rubified soils (G1); steep hillslopes in conglomerate, with shallow and poorly developed soils (H1); low-gradient hillslopes in conglomerate, with strongly decarbonated, rubified soils with evidence of clay illuviation (H2); steep hillslopes in sandstone, with moderately deep and developed soils (H3); low-gradient hillslopes in marls and siltite, with moderately deep and developed soils (H4); long and steep mountain slopes in massive and hard limestone, with shallow and poorly developed soils (V1); long and steep mountain slopes in well-stratified, moderately resistant limestone, with moderately deep and leached soils with clay illuviation (V2).

The model shows zones with low values near to 0 Mg ha^−1^ year^−1^ mainly in the gravelly alluvial plain, grassland, forests or slope near 0° degrees; conversely, the highest erosion rate values (more of 400 Mg ha^−1^ year^−1^) are distributed on steepest slopes, mostly on bare soil areas. Generally, erosion rate with bigger intensity (>40 Mg ha^−1^ year^−1^) is clustered on long and steep slopes, characterized by intensive agricultural activities (Fig 5). Here, specific land use determines different effects on potential soil erosion rate: olive groves (67.4 Mg ha^−1^ year^−1^), vineyards (59.8 Mg ha^−1^ year^−1^), “other permanent crops” (46.2 Mg ha^−1^ year^−1^). As expected, soil erosion rate is more intensive on hilly landscapes, characterized by intensive vineyards cropland, as it shown in Figures 4b, 4c and 5.

**Fig 5.**
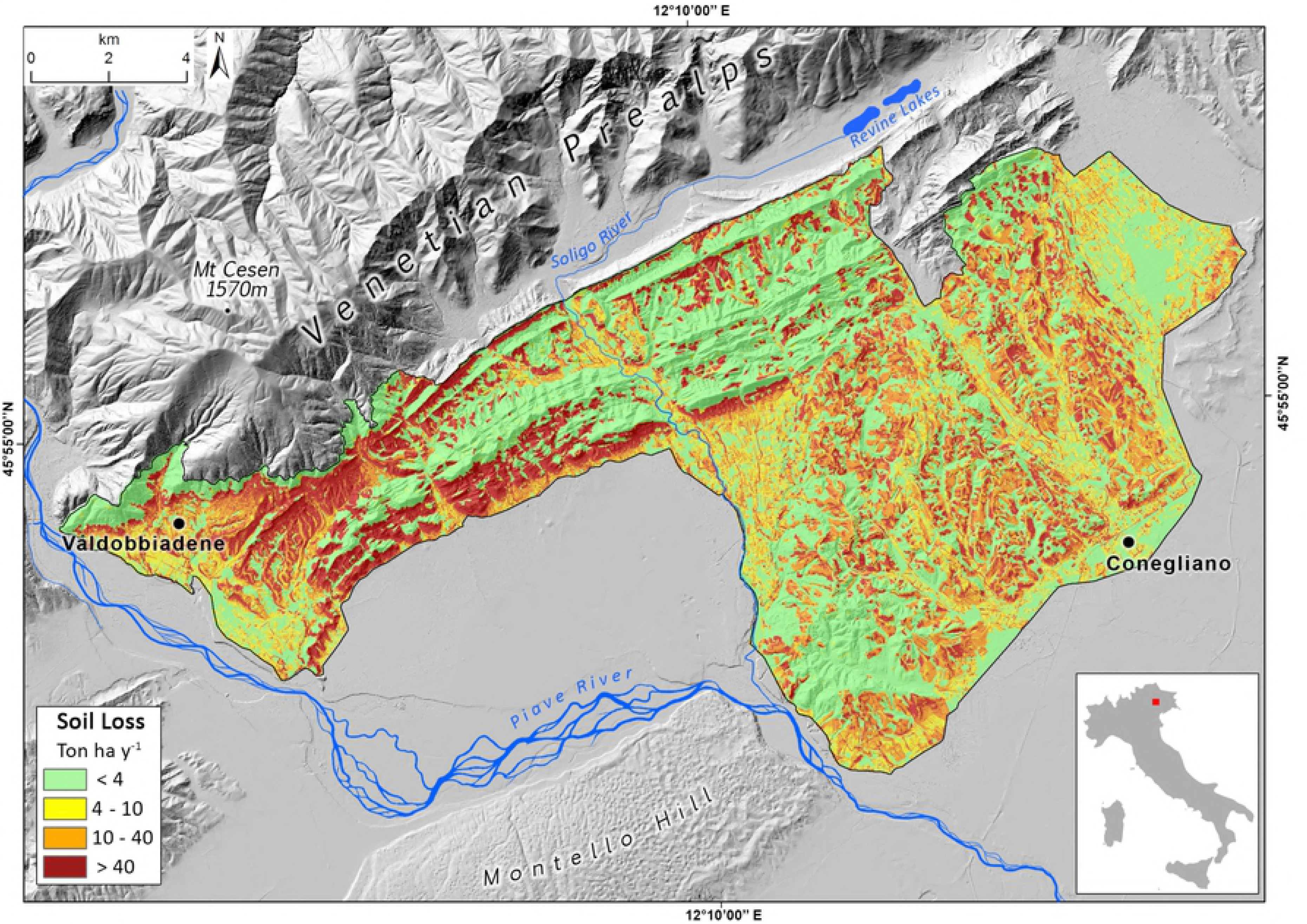
Map of soil erosion rate in the Prosecco DOCG area represented in four classes.

If we consider the total soil erosion modelled for all the Prosecco DOCG area, the RUSLE analysis shows that vineyards contribute for 400,000 Mg year^−1^, which contributes to the 74% of all the erosion potential in the whole area (Fig 4b). Therefore, if the average of the declared wine production in the last years is more than 90 M bottles, a single bottle of Prosecco DOCG sparkling wine embodies a “soil footprint” on the territory of about 4.4 kg year^−1^.

In the study area, soil erosion seems to be potentially higher in soil unit systems H4 and H1 which represent soils in hilly landscapes (Fig 4c). In fact, more than the 58% of soil erosion potential is focused in the hilly sector of the Prosecco DOCG area. Furthermore, H1 plus H4 represent more than 51% of all territory surface (Fig 4d).

As it is illustrated in figure 4e, if we consider only soil erosion potential expressed by hectares (Mg ha^−1^ year^−1^) the soil system showing the highest values is P2, losing more than twice (about 9 Mg ha^−1^ year^−1^) in respect to H4 (only 4 Mg ha^−1^ year^−1^). However, P2 soil system covers only 600 ha and its contribution to total soil erosion is very low. Soil systems are more susceptible to erosion than P2, M3 and C2; however, with the exception of C2, they have limited extension in the study area (Fig 4d). C2 represents recent soils not decarbonated, which are found in the valley bottom. It is the third soil system unit for surface. It is an example of an high erosion potential zone and it could be a risk area.

In the Prosecco wine district, the highest soil erosion potential is localized within the town of “Valdobbiadene”, which covers about the 12% of total Prosecco DOCG surface. Here, soil erosion risk is about 100,000 Mg year^−1^, corresponding to the 18% of total soil erosion estimation in the Prosecco DOCG area, followed by Farra di Soligo (94,000 Mg year^−1^, the 17% of the area) and Conegliano (61,000 Mg year^−1^, the 11% of the area). Municipalities with the highest average rates of soil erosion are: Farra di Soligo (more than 60 Mg ha^−1^ year^−1^), Vidor (57 Mg ha^−1^ year^−1^), and Miane (38 Mg ha^−1^ year^−1^).

### Alternative sustainable land-management scenarios

In the first simulated sustainable scenario, 5 m grassed buffer filter-strips modelled around rivers and streams (197 ha) show a total erosion potential of 502,623 Mg year^−1^, representing a reduction of the 7.8% of soil loss. In the second scenario, a reduction of 55,715 Mg year^−1^ (10.1%) in soil erosion rate was obtained by simulating a mitigation measure of 3,5 m of hedgerows around vineyards, accounting for a total of 645 ha. An important reduction in soil erosion is obtained by summarizing the mitigation effects of buffer filter-strips, both around the river networks and vineyards plots: soil loss erosion may be reduced of 14.8%, which corresponds to 80,458 Mg year^−1^ of soil preserved (Fig 3f).

However, the most sustainable scenario in our analyses is represented by simulating a best practice of leaving the 100% grass cover of vine inter-rows. In this case, the total erosion in the Prosecco DOCG area would be reduced to 275,140 Mg year^−1^, saving about the 50% of soil loss. In vineyards a general decrease of almost 3 times (from 400,000 to 135,161 Mg year^−1^) is also demonstrated, reducing on average the erosion rate from 59.8 to 19.2 Mg ha^−1^ year^−1^. In this more sustainable scenario total erosion related to vine production in the all Prosecco DOCG area is reduced from 70% to 49%.

## Discussion

### Sparkling earth for Prosecco wine

According to estimation by Cerdan et al. (2010), the mean erosion rate in Italy is 2.3 Mg ha^−1^ year [13]; in our study we found that in the Prosecco DOCG area the erosion rate modelled is 25.4 Mg ha year^−1^, a magnitude of erosion rate which is 11 times higher than the Italian average. We found soil erosion rate 3.8 times higher than in the Aosta valley vineyards (NW Italy), in a similar morphological context and agricultural management practices [49]. According to Aiello et al. (2015) [51], which computed a modified RUSLE model for complex terrain (RUSLE3D) along a highly-erodible hilly landscape in Basilicata (Southern Italy), the mean annual soil erosion in the Bradano basin is 31.80 Mg ha^−1^ year^−1^, which is 1.8 lower than values we found in the Prosecco DOCG area.

This study confirmed the key role of vineyards in soil erosion processes, contributing to the highest values (>40 Mg ha^−1^ year^−1^), mainly clustered in the hilly areas, especially on steep slopes (Figs. 5 and 6). This is the case in the areas of Valdobbiadene and Farra di Soligo (Province of Treviso) which account for the 18% and the 17% of the total soil erosion in the Prosecco DOCG. The average erosion rate we modelled in Prosecco vineyards is 59.8 Mg ha^−1^ year^−1^, which is 40 times greater than the upper limit of tolerable soil erosion threshold defined for Europe by Verhejen et al. (2009) [11]. Similar results based on the RUSLE model were found by Prosdocimi et al (2016) in the Lierza river basin of the Prosecco DOCG area [20]. Other results performed in experimental plots in sloping vineyards in Germany (Mosel Valley), Eastern Spain (Les Alcusses Valley), and Southwestern Sicily (Agrigento) confirmed soil erosion under conventional land management range from 19 to 102 Mg ha^−1^ year^−1^ [21,24,62].

**Fig 6.**
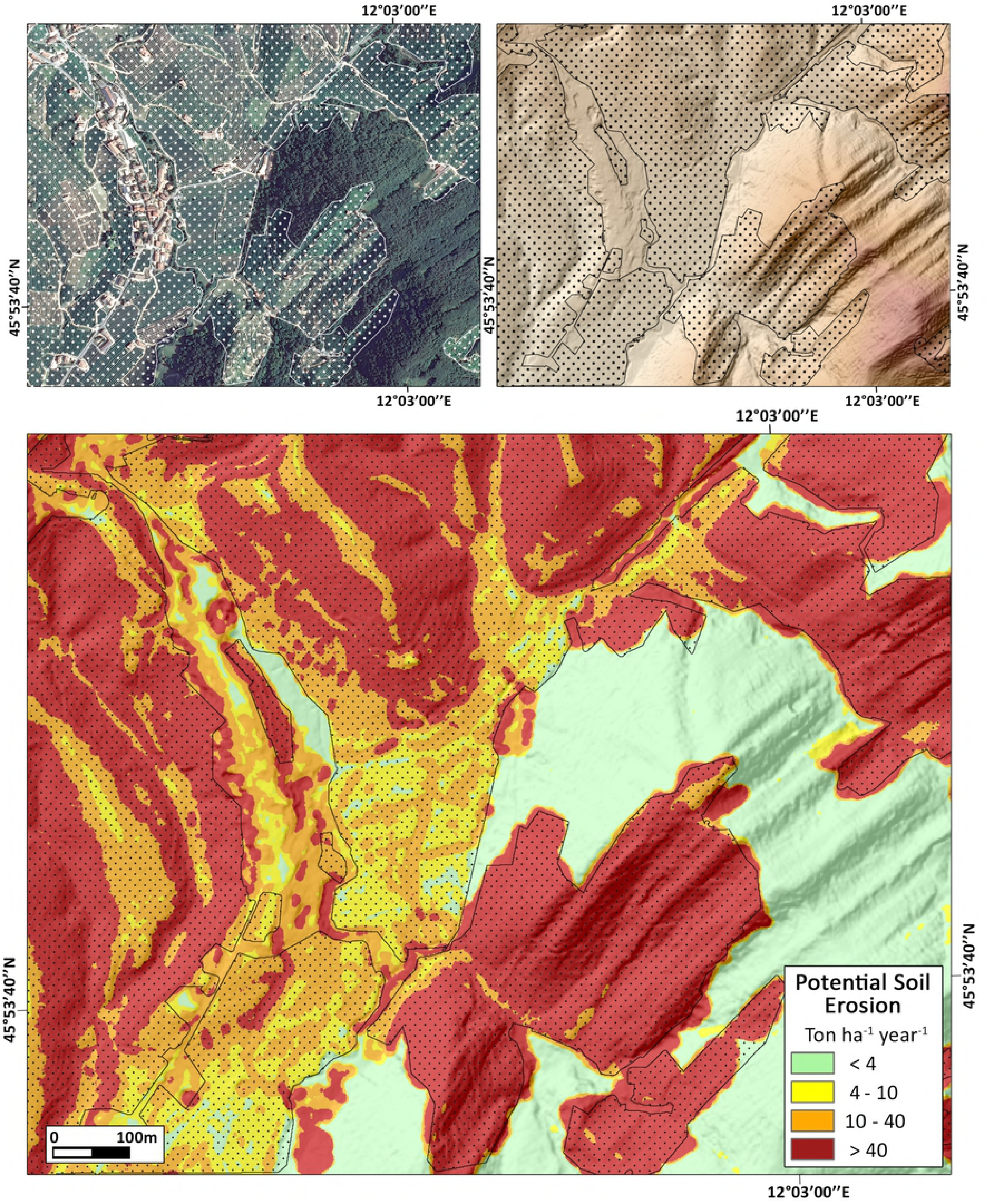
Sample area of S. Stefano di Barbozza (Valdobbiadene Municipality). Upper left: Aerial photo of the village and its surrounding. Upper right: DTM of the same area. Lower inset: map of potential soil loss from RUSLE modelling. Polygons with hatching indicate vineyards.

It is worth noting that we performed the most conservative scenario for soil erosion in a conventional and a chemically weeded vineyard, by using a C standard value of 0.12 for vineyards for RUSLE analyses, not taking into account direct effects of the modern agrarian-hydraulic layouts, where terrain morphology and drainage system are strongly modified by vineyards row setup along the steepest slope to facilitate agricultural operations.

In fact, particular concern is presently given to new vine plantations which are increasing on hillslopes of the Prosecco DOCG area. As it is widely documented they trigger to extreme erosion rates due to drastic changes in soil physical properties through heavy levelling operations, deep ploughing, trampling, and down-slope orientation of vine-rows. Moreover, inter-rows maintenance with bare soil or soil scarcely vegetated by grass cover (5-30%), result in heavy runoff and, therefore, increasing soil erosion rates [49,63]. Different studies highlight that new vine plantations strongly contributes to high erosion risk by increasing rates up to 30 times higher than the upper threshold for tolerable erosion suggested in Europe [11,21,64,65].

High soil erosion rate may exacerbate in-site effects significantly affecting crop production in soil quality and fertility reduction by decrease in nutrients and organic matter [11,12,48,66]. Moreover, considering the emerging climate change scenarios in Mediterranean regions, an increase in frequency of extreme rainfall events in spring and autumn, especially after dry periods, may amplify off-site impacts on steep slopes by soil water erosion and heavy runoff [67–70]. Off-site impacts are related to non-point source pollution from agricultural fields: pesticide and fertilizers runoff into stream and river network, contamination of groundwater resources, and air pollution by emission of greenhouse gasses such as CO_2_, CH_4_ and N_2_O [9,11,71]. Moreover, high erosion rates may affect slope stability, amplifying hydrogeological risk [72,73]. This suggest that in mid-long-term degradation in ecosystem functioning could strongly affect agricultural productivity by drastic reduction in nutrients, organic matter, water capacity and biota.

As it is widely recognized soils are the base of a wide-set of ecosystem good and services which are fundamental for human needs: food production, drinking water quality, water purification, hydrogeological risk control, biodiversity and carbon stock shrinkage. Hence, erosion processes directly lead to degradation and loss of ecosystem services, undermining soil sustainability as recognized both in the seven soil functions defined by the European Commission (2006) and the land-related 2030 UN Sustainable Development Goals [40,41]. Moreover, the European Union brought this issue into the current environmental policy agenda by including soil erosion among the eight soil threats listed within the Soil Thematic Strategy of the European Commission (EC, 2006) and in different policy developments such as the Common Agricultural Policy (CAP), Europe 2020, and the 7th Environmental Action Programme [5].

### Walking pathways by nature-based agricultural practices

The four simulations of sustainable land management scenarios which are included in the GAEC standards show that minor variations in land use could significantly change the total soil loss in the study area. The combination of 5 m grassed buffer filter-strips together with 3,5 m of hedgerows around vineyards potentially preserve 80,458 Mg year^−1^ of soil. Moreover, as reported in an experimental trial performed in the Prosecco DOCG area, hedgerows represent also an effective mitigation measure to reduce up the 95-98% the spray drift effect of pesticide from vineyards [74].

On the other hand, the forth sustainable scenario which simulate to shift from intensive herbicide application to a 100% inter-row vine grass cover during Winter time demonstrates the potential effectiveness of this nature-based solution, by reducing soil erosion rate within Prosecco vineyards of 66.2%. Results of the simulated mitigation effect are not so different from those derived from experimental measures at field-scale in Sicilian vineyards, where different cover crops sowed in vine inter-rows reduced soil erosion by 68% compared with conventional land management [24]. Similar results were also found in Spain through different erosion measures under simulated rainfall where cover crop of *Secale sp.* and *Brachipodium sp*. showed a significant reduction in soil erosion [75]; another immediate effectiveness of field-scale mitigation measures is represented by the use of barley straw mulch in Mediterranean vineyards which reduces the median erosion from 2.81 to 0.63 80,458 Mg year^−1^ [76]. The most effective mitigation effects at field-scale seems to be the use of straw as mulch together with no-tillage strategy which can reduce soil erosion rate of two orders of magnitude [13].

In intensive vineyard croplands such as the Prosecco DOCG, mitigation measures and best management practices should be adopted in the GAEC framework, which also provide economic incentives to farmers which implement in-site measures like hedgerows and/or grassed buffer filter strips, dry-stone walls terraces, contour farming, and strip cropping to control soil erosion processes in vineyard cropland and to reduce off-site impacts.

## Conclusions

The RUSLE model was applied to estimate the total soil erosion in the Prosecco DOCG, by identifying the most critical zones and by simulating alternative nature-based land management scenarios at landscape scale. This study confirmed the key role of Prosecco vineyards in increasing soil erosion processes, by contributing to the 74% of total erosion in the DOCG area with in-field rate 40 times greater than the upper limit of tolerable soil erosion threshold defined for Europe.

This suggest that i) in mid-long-term period degradation in ecosystem functioning could strongly affect agricultural productivity by drastic reduction in nutrients, organic matter, water capacity and biota; ii) off-site effects such as leaching of agricultural pollutants and erosion risk may affect at multiple scale in the territory.

Using a RUSLE GIS-based approach we modelled a “soil footprint” for producing a single bottle of Prosecco DOCG sparkling wine, which currently “drinks” about 4.4 kg of soil every year.

On the other hand, nature-based agricultural practices showed relevant decrease in soil erosion and they are strongly recommended in the DOCG area, especially where erosion rate is critical.

Our study suggests that in the Prosecco DOCG an integrated soil erosion monitor system is needed area, combining field measures with spatial analyses at territory scale.

## Acknowledments

We would thank the Second Level Master in GIScience and UAV (ICEA Department, University of Padua) and GIS lab of the Geography Section (DiSSGeA Department, University of Padua) for the technical support.

## References

1. FAO. FAOSTAT. 2017 [cited 14 Dec 2018]. Available: http://www.fao.org/faostat/en/#data/EL

2. FAO. The State of Food and Agriculture. Livestock in the Balance. 2016. doi:ISBN: 978-92-5-107671-2 I

3. Stallman HR. Ecosystem services in agriculture: Determining suitability for provision by collective management. Ecol Econ. Elsevier B.V.; 2011;71: 131–139. doi:10.1016/j.ecolecon.2011.08.016

4. Tilman D, Cassman KG, Matson PA, Naylor R, Polasky S. Nature01017. 2002;418. doi:10.1080/11263508809430602

5. Campbell BM, Beare DJ, Bennett EM, Hall-Spencer JM, Ingram JSI, Jaramillo F, et al. Agriculture production as a major driver of the earth system exceeding planetary boundaries. Ecol Soc. 2017;22. doi:10.5751/ES-09595-220408

6. Montgomery DR. Soil erosion and agricultural sustainability. Proc Natl Acad Sci U S A. 2007;104: 13268–72. doi:10.1073/pnas.0611508104

7. Gabriels D, Cornelis WM. Human-Induced Land Degradation. Land Use, Land Cover And Soil Sciences. 2009. pp. 131–143.

8. Parras-AlcÃ¡ntara L, Lozano-GarcÃ-a B, Keesstra S, CerdÃ A, Brevik EC. Long-term effects of soil management on ecosystem services and soil loss estimation in olive grove top soils. Sci Total Environ. Elsevier B.V.; 2016;571: 498–506. doi:10.1016/j.scitotenv.2016.07.016

9. Jahun BG, Ibrahim R, Dlamini NS, Musa SM. Review of Soil Erosion Assessment using RUSLE Model and GIS. J Biol Agric Healthc. 2015;5: 36–47.

10. Panagos P, Borrelli P, Meusburger K, van der Zanden EH, Poesen J, Alewell C. Modelling the effect of support practices (P-factor) on the reduction of soil erosion by water at European scale. Environ Sci Policy. Elsevier Ltd; 2015;51: 23–34. doi:10.1016/j.envsci.2015.03.012

11. Verheijen FGA, Jones RJA, Rickson RJ, Smith CJ. Tolerable versus actual soil erosion rates in Europe. Earth-Science Rev. Elsevier B.V.; 2009;94: 23–38. doi:10.1016/j.earscirev.2009.02.003

12. Panagos P, Borrelli P, Poesen J, Ballabio C, Lugato E, Meusburger K, et al. The new assessment of soil loss by water erosion in Europe. Environ Sci Policy. Elsevier Ltd; 2015;54: 438–447. doi:10.1016/j.envsci.2015.08.012

13. Cerdan O, Govers G, Le Bissonnais Y, Van Oost K, Poesen J, Saby N, et al. Rates and spatial variations of soil erosion in Europe: A study based on erosion plot data. Geomorphology. Elsevier B.V.; 2010;122: 167–177. doi:10.1016/j.geomorph.2010.06.011

14. Gómez J, Infante-Amate J, de Molina M, Vanwalleghem T, Taguas E, Lorite I. Olive Cultivation, its Impact on Soil Erosion and its Progression into Yield Impacts in Southern Spain in the Past as a Key to a Future of Increasing Climate Uncertainty. Agriculture. 2014;4: 170–198. doi:10.3390/agriculture4020170

15. Ramos MI, Feito FR, Gil AJ, Cubillas JJ. A study of spatial variability of soil loss with high resolution DEMs: A case study of a sloping olive grove in southern Spain. Geoderma. Elsevier B.V.; 2008;148: 1–12. doi:10.1016/j.geoderma.2008.08.015

16. Keesstra S, Pereira P, Novara A, Brevik EC, Azorin-Molina C, Parras-Alcántara L, et al. Effects of soil management techniques on soil water erosion in apricot orchards. Sci Total Environ. Elsevier B.V.; 2016;551–552: 357–366. doi:10.1016/j.scitotenv.2016.01.182

17. Cerdà A, Giménez-Morera A, Bodì MB. Soil and water losses from new citrus orchards growing on sloped soils in the western Mediterranean basin. Earth Surf Process Landforms. 2009;34: 1822–1830. doi:10.1002/esp.1889

18. Aurand JM. State of the Vitiviniculture World Market. 38th OIV World Congress of vine and wine. 2015. pp. 1–14.

19. OIV Organisation Internationale de la Vigne et du Vin. State of the vitiviniculture world market. 2018 pp. 1–14. Available: http://www.oiv.int/en/technical-standards-and-documents/statistical-analysis/state-of-vitiviniculture

20. Prosdocimi M, Cerdà A, Tarolli P. Soil water erosion on Mediterranean vineyards: A review. Catena. Elsevier B.V.; 2016;141: 1–21. doi:10.1016/j.catena.2016.02.010

21. Cerdà A, Keesstra SD, Rodrigo-Comino J, Novara A, Pereira P, Brevik E, et al. Runoff initiation, soil detachment and connectivity are enhanced as a consequence of vineyards plantations. J Environ Manage. 2017;202: 268–275. doi:10.1016/j.jenvman.2017.07.036

22. Rodrigo Comino J, Iserloh T, Morvan X, Malam Issa O, Naisse C, Keesstra S, et al. Soil Erosion Processes in European Vineyards: A Qualitative Comparison of Rainfall Simulation Measurements in Germany, Spain and France. Hydrology. 2016;3: 6. doi:10.3390/hydrology3010006

23. Mariani A, Pomarici E, Boatto V. The international wine trade: Recent trends and critical issues. Wine Econ Policy. 2012;1: 24–40. doi:10.1016/j.wep.2012.10.001

24. Novara A, Gristina L, Saladino SS, Santoro A, Cerdà A. Soil erosion assessment on tillage and alternative soil managements in a Sicilian vineyard. Soil Tillage Res. 2011;117: 140–147. doi:10.1016/j.still.2011.09.007

25. Norbiato D, Borga M, Dinale R. Flash flood warning in ungauged basins by use of the flash flood guidance and model-based runoff thresholds. Meteorol Appl. 2009;16: 65–75. doi:10.1002/met.126

26. Cerdá A. Soil aggregate stability in three Mediterranean environments. Soil Technol. 1996;9: 133–140.

27. Lasanta T, Arnáez J, Oserín M, Ortigosa LM, Study AC, Viejo C, et al. Marginal Lands and Erosion in Terraced Fields in the Mediterranean Mountains Marginal Lands and Erosion in Terraced Fields in the Mediterranean Mountains. 2001;21: 69–76.

28. Rodrigo Comino J, Iserloh T, Lassu T, Cerdà A, Keesstra SD, Prosdocimi M, et al. Quantitative comparison of initial soil erosion processes and runoff generation in Spanish and German vineyards. Sci Total Environ. Elsevier B.V.; 2016;565: 1165–1174. doi:10.1016/j.scitotenv.2016.05.163

29. Santisteban LM De, Casalí J, López JJ. Assessing soil erosion rates in cultivated areas of Navarre (Spain). 2006;506: 487–506. doi:10.1002/esp.1281

30. Pomarici E. Recent trends in the international wine market and arising research questions. Wine Econ Policy. 2016;5: 1–3. doi:10.1016/j.wep.2016.06.001

31. Varotto M, Tress M. Paesaggi in movimento: il difficile equilibrio tra permanenze e trasformazioni in Valsana. Esercizi di paesaggio. Regione del Veneto, Direzione Urbanistica e Paesaggio; 2011. pp. 111–124.

32. Visentin F, Vallerani F. A Countryside to Sip : Venice Inland and the Prosecco’s Uneasy Relationship with Wine Tourism and Rural Exploitation. Sustainability. 2018; 2195. doi:10.3390/su10072195

33. ISPRA Istituto Superiore per la Protezione e la Ricerca Ambientale. Consumo di suolo, dinamiche territoriali e servizi ecosistemici. ISPRA. Roma; 2018.

34. Tomasi D, Gaiotti F, Jones G V. The power of the terroir: The case study of prosecco wine. The Power of the Terroir: The Case Study of Prosecco Wine. 2013. doi:10.1007/978-3-0348-0628-2_1

35. Esteri R. Le colline del Prosecco non entrano nel Patrimonio dell’umanità Unesco. In: Il Corriere.it. 2018 pp. 18–21.

36. De Marchi A. Colline del Prosecco patrimonio Unesco: arriva il primo “no.” In: La Tribuna di Treviso. 2018 pp. 1–12.

37. Onofri L, Boatto V, Bianco AD. Who likes it “sparkling”? An empirical analysis of Prosecco consumers’ profile. 2015; doi:10.1186/s40100-014-0026-x

38. Vinitaly. Il Prosecco Docg cresce più in valore che in volum - Vinitaly [Internet]. [cited 14 Jul 2017]. Available: http://www.vinitaly.com/it/news/wine-news/il-prosecco-docg-cresce-piu-in-valore-che-in-volum/

39. Emanuele Scarci. La crescita record del Prosecco. Il Sole 24 ORE. 21 Dec 2016. Available: http://www.ilsole24ore.com/art/impresa-e-territori/2016-12-20/la-crescita-record-prosecco-143556.shtml?uuid=AD994JHC&refresh_ce=1. Accessed 13 Jul 2017.

40. Caputo R, Poli ME, Zanferrari A. Neogene–Quaternary tectonic stratigraphy of the eastern Southern Alps, NE Italy. J Struct Geol. 2010;32: 1009–1027. doi:10.1016/j.jsg.2010.06.004

41. Doglioni C. Thrust tectonics examples from the Venetian Alps. Stud Geol Camerti, Vol Spec 117-129. 1990; 117–129. Available: http://www.academia.edu/5531251/Thrust_tectonics_examples_from_the_Venetian_Alps

42. Venzo S. I depositi quaternari e del neogene superiore nella bassa valle del Piave da Quero al Montello e del paleopiave nella valle del Soligo (Treviso) [Internet]. Padova: Istituti di geologia e mineralogia dell’Università de Padova; 1977. Available: http://www.worldcat.org/title/depositi-quaternari-e-del-neogene-superiore-nella-bassa-valle-del-piave-da-quero-al-montello-e-del-paleopiave-nella-valle-del-soligo-treviso/oclc/66003587

43. Carton A, Bondesan A, Fontana A, Meneghel M, Miola A, Mozzi P, et al. Geomorphological evolution and sediment transfer in the Piave River system (northeastern Italy) since the Last Glacial Maximum. Géomorphologie Reli Process Environ. Groupe français de géomorphologie; 2009;15: 155–174. doi:10.4000/geomorphologie.7639

44. Agenzia Regionale per la Prevenzione e protezione ambientale del Veneto. ARPAV. La Carta dei Suoli della Provincia di Treviso. Treviso: Provincia di Treviso; 2008.

45. Renard KG, Foster GR, Weesies Gienn A, Porter Jeffrey p. RUSLE: Revised universal soil loss equation. J Soil Water Conserv. Soil Conservation Society of America]; 1991;46: 30–33. Available: http://www.jswconline.org/content/46/1/30.extract

46. Wischmeier W, Smith DD, Wischmer WH, Smith DD. Predicting rainfall erosion losses: a guide to conservation planning. US Dep Agric Handb No 537. 1978; 1–69. doi:10.1029/TR039i002p00285

47. Ashiagbor G, Forkuo EK, Laari P, Aabeyir R. Modeling Soil Erosion Using Rusle and Gis Tools. Int J Remote Sens Geosci. 2013;2. Available: www.ijrsg.com

48. Prosdocimi M, Cerdà A, Tarolli P. Soil water erosion on Mediterranean vineyards: A review. Catena. Elsevier B.V.; 2016;141: 1–21. doi:10.1016/j.catena.2016.02.010

49. Biddoccu M, Zecca O, Audisio C, Godone F, Barmaz A, Cavallo E. Assessment of Long-Term Soil Erosion in a Mountain Vineyard, Aosta Valley (NW Italy). L Degrad Dev. 2017; doi:10.1002/ldr.2657

50. Biddoccu M, Ferraris S, Cavallo E, Opsi F, Previati M, Canone D. Hillslope Vineyard Rainfall-Runoff Measurements in Relation to Soil Infiltration and Water Content. Procedia Environ Sci. 2013;19: 351–360. doi:10.1016/j.proenv.2013.06.040

51. Aiello A, Adamo M, Canora F. Remote sensing and GIS to assess soil erosion with RUSLE3D and USPED at river basin scale in southern Italy. Catena. Elsevier B.V.; 2015;131: 174–185. doi:10.1016/j.catena.2015.04.003

52. Lal R. Soil erosion in the tropics?: principles and management [Internet]. McGraw-Hill; 1990. Available: http://agris.fao.org/agris-search/search.do?recordID=US9120604

53. Regione Veneto. Carta di Copertura del Suolo della Regione del Veneto. GSE Land–Urban Atlas [Internet]. Regione del Veneto; 2012. Available: http://idt.regione.veneto.it/app/metacatalog/getMetadata/?id=551

54. EEA. The revised and supplemented Corine land cover nomenclature. EEA Tech Rep No 40. 2000; 110.

55. Agenzia Regionale per la Prevenzione e protezione ambientale del Veneto. ARPAV. Valutazione del rischio di erosione per la regione Veneto [Internet]. 2008. Available: http://www.arpa.veneto.it/temi-ambientali/suolo/file-e-allegati/documenti/minacce-di-degradazione/Rapportofinale_erosione_ARPAV3.pdf

56. Bazzoffi P. Erosione del suolo e sviluppo rurale?: fondamenti e manualistica per la valutazione agroambientale [Internet]. Edagricole; 2007. Available: https://www.libreriauniversitaria.it/erosione-suolo-sviluppo-rurale-sostenibile/libro/9788850652280

57. Bazzoffi P. Soil erosion tolerance and water runoff control: Minimum environmental standards. Reg Environ Chang. 2009;9: 169–179. doi:10.1007/s10113-008-0046-8

58. D. K. McCool DK, L. C. Brown LC, G. R. Foster GR, C. K. Mutchler CK, L. D. Meyer LD. Revised Slope Steepness Factor for the Universal Soil Loss Equation. Trans ASAE. American Society of Agricultural and Biological Engineers; 1987;30: 1387–1396. doi:10.13031/2013.30576

59. Blanos R, De cillia C, Paganini P, Pavan A, Sterzai P, Pietrapertosa C, et al. Rilievo lidar ed iperspettrale della provincia di Treviso. 2009.

60. Torri D, Poesen J, Borselli L. Predictability and uncertainty of the soil erodibility factor using a global dataset. CATENA. 1997;31: 1–22. doi:10.1016/S0341-8162(97)00036-2

61. Bazzoffi P. Soil erosion tolerance and water runoff control: minimum environmental standards. Reg Environ Chang. Springer-Verlag; 2009;9: 169–179. doi:10.1007/s10113-008-0046-8

62. Rodrigo Comino J, Quiquerez A, Follain S, Raclot D, Le Bissonnais Y, Casalí J, et al. Soil erosion in sloping vineyards assessed by using botanical indicators and sediment collectors in the Ruwer-Mosel valley. Agric Ecosyst Environ. Elsevier B.V.; 2016;233: 158–170. doi:10.1016/j.agee.2016.09.009

63. Tropeano D. Rate of soil erosion processes on vineyards in central Piedmont (NW Italy). Earth Surf Process Landforms. John Wiley & Sons, Ltd; 1984;9: 253–266. doi:10.1002/esp.3290090305

64. Rodrigo-Comino J, Seeger M, Senciales JM, Ruiz-Sinoga JD, Ries JB. Variación espacio-temporal de los procesos hidrológicos del suelo en viñedos con elevadas pendientes (valle del ruwer-mosela, alemania). Cuad Investig Geogr. 2016;42: 281–306. doi:10.18172/cig.2934

65. Kirchhoff M, Seeger M, Ries JB. Soil Erosion in Sloping Vineyards Under Conventional and Organic Land Use. 2017;43: 119–140.

66. Maetens W, Vanmaercke M, Poesen J, Jankauskas B, Jankauskiene G, Ionita I. Effects of land use on annual runoff and soil loss in Europe and the Mediterranean. Prog Phys Geogr. SAGE PublicationsSage UK: London, England; 2012;36: 599–653. doi:10.1177/0309133312451303

67. Capolongo D, Pennetta L, Piccarreta M, Fallacara G, Boenzi F. Spatial and temporal variations in soil erosion and deposition due to land-levelling in a semi-arid area of Basilicata (Southern Italy). Earth Surf Process Landforms. John Wiley & Sons, Ltd.; 2008;33: 364–379. doi:10.1002/esp.1560

68. Borga M, Anagnostou EN, Blöschl G, Creutin J-D. Flash flood forecasting, warning and risk management: the HYDRATE project. Environ Sci Policy. 2011;14: 834–844. doi:10.1016/j.envsci.2011.05.017

69. Sofia G, Roder G, Dalla Fontana G, Tarolli P. Flood dynamics in urbanised landscapes: 100 years of climate and humans’ interaction. Sci Rep. 2017;7: 40527. doi:10.1038/srep40527

70. Zollo AL, Rillo V, Bucchignani E, Montesarchio M, Mercogliano P. Extreme temperature and precipitation events over Italy: assessment of high-resolution simulations with COSMO-CLM and future scenarios. Int J Climatol. John Wiley & Sons, Ltd; 2016;36: 987–1004. doi:10.1002/joc.4401

71. Lollino G, Manconi A, Clague J, Shan W, Chiarle M. Effects of Soil Management on Long-Term Runoff and Soil Erosion Rates in Sloping Vineyards. Engineering Geology for Society and Territory. 2015. pp. 1–572. doi:10.1007/978-3-319-09300-0

72. Onori F, De Bonis P, Grauso S. Soil erosion prediction at the basin scale using the revised universal soil loss equation (RUSLE) in a catchment of Sicily (southern Italy). Environ Geol. 2006;50: 1129–1140. doi:10.1007/s00254-006-0286-1

73. Evans R, Boardman J. The new assessment of soil loss by water erosion in Europe. Panagos P. et al., 2015 Environmental Science & Policy 54, 438-447-A response. Environ Sci Policy. Elsevier Ltd; 2016;58: 11–15. doi:10.1016/j.envsci.2015.12.013

74. Otto S, Loddo D, Baldoin C, Zanin G. Spray drift reduction techniques for vineyards in fragmented landscapes. J Environ Manage. Elsevier Ltd; 2015;162: 290 298. doi:10.1016/j.jenvman.2015.07.060

75. Bienes R, Mun G, Marques MJ, Garci S. SOIL CONSERVATION BENEATH GRASS COVER IN HILLSIDE VINEYARDS UNDER MEDITERRANEAN CLIMATIC CONDITIONS (MADRID, SPAIN). 2010;131: 122–131.

76. Prosdocimi M, Jordán A, Tarolli P, Keesstra S, Novara A, Cerdà A. The immediate effectiveness of barley straw mulch in reducing soil erodibility and surface runoff generation in Mediterranean vineyards. Sci Total Environ. Elsevier B.V.; 2016;547: 323–330. doi:10.1016/j.scitotenv.2015.12.076

